# Structural and biochemical analysis of OrfG: the VirB8-like component of the integrative and conjugative element ICE*St3* from *Streptococcus thermophilus*

**DOI:** 10.1101/2020.11.26.395491

**Authors:** Julien Cappele, Abbas Mohamad-Ali, Nathalie Leblond-Bourget, Sandrine Mathiot, Tiphaine Dhalleine, Sophie Payot, Martin Savko, Claude Didierjean, Frédérique Favier, Badreddine Douzi

## Abstract

Conjugative transfer is a major threat to global health since it contributes to the spread of antibiotic resistance genes and virulence factors among commensal and pathogenic bacteria. To allow their transfer, mobile genetic elements including Integrative and Conjugative Elements (ICEs) use a specialized conjugative apparatus related to Type IV secretion systems (Conj-T4SS). Therefore, Conj-T4SSs are excellent targets for strategies that aim to limit the spread of antibiotic resistance. In this study, we combined structural, biochemical and biophysical approaches to study OrfG, a protein that belongs to Conj-T4SS of ICE*St3* from *Streptococcus thermophilus*. Structural analysis of OrfG by X-ray crystallography revealed that OrfG central domain is similar to VirB8-like proteins but displays a different quaternary structure in the crystal. To understand, at a structural level, the common and the diverse features between VirB8-like proteins from both Gram-negative and -positive bacteria, we used an *in silico* structural alignment method that allowed us to identify different structural classes of VirB8-like proteins. Biochemical and biophysical characterizations of purified OrfG soluble domain and its central and C-terminal subdomains indicated that they are mainly monomeric in solution but able to form an unprecedented 6-mer oligomers. Our study provides new insights into the structural and assembly mode of VirB8-like proteins, a component essential for conjugative transfer and improves our understanding on these under-examined bacterial nanomachines.

## Introduction

Conjugation of mobile genetic elements constitutes one of the major mechanisms of horizontal gene transfer [1]. Besides their crucial role in prokaryotic evolution and adaptation, conjugative elements including plasmids and integrative conjugative elements (ICEs) constitute the most important vectors of antibiotic resistance spreading among bacteria. Conjugative transfer is ensured by dynamic multi-subunit cell-envelope spanning nanomachines called conjugative type IV secretion systems (Conj-T4SSs) encoded by conjugative elements.

Based on the well-studied archetypal VirB/D4 Conj-T4SS system of the Ti conjugative plasmid from the Gram-negative bacterium *Agrobacterium tumefaciens*, it is commonly assumed that Vir subunits can be grouped into 3 classes [2–5]. One encloses the energetic components VirB4, VirB11 and VirD4. VirB4 and VirB11 belong to traffic ATPases and are involved in the functioning of T4SS apparatus and in pili biogenesis whereas the coupling protein VirD4 (TC4P) is involved in DNA substrate recognition. The second class corresponds to the inner membrane platform, which is composed of the membrane proteins VirB3, VirB6 and VirB8. The inner membrane platform serves as a docking station for DNA substrate and constitutes the inner membrane channel. The third class forms the outer membrane core complex. It is composed of VirB7, VirB9 and VirB10 proteins. The outer membrane core complex forms a conducting channel that traverses the periplasmic space and the outer membrane barrier, establishing a continuity for the inner membrane channel. Other proteins are involved in the conjugative process. These proteins include the transglycosylase VirB1 that is important for the establishment of the T4SS apparatus within the peptidoglycan layer in the periplasmic space and of the extracellular pili involved in DNA transfer or in host cell attachment. Conjugative pili are predominantly composed by the VirB2 pilin and are tipped by the VirB5 adhesin [5, 6]. DNA transfer process through Conj-T4SS requires DNA recruitment and delivery to the T4SS apparatus. Initial steps of DNA recruitment are conducted by the relaxosome complex constituted by the relaxase and accessory factors that recognise and bind specifically to the origin of transfer *oriT* sequence located in the conjugative element. The nick of the *oriT* sequence is catalysed by the relaxase that binds covalently to the nascent 5’ end of the DNA strand to be transferred (T-strand). Straight after, the relaxase-T-strand nucleoprotein docks to T4CP, which consequently unfolds the relaxase and ensures the transfer of the nucleoprotein to the Conj-T4SS channel. Once transferred to the recipient cell, relaxase catalyses the recircularization of the T-strand followed by the second strand synthesis [7].

Conj-T4SSs in Gram-positive bacteria are more puzzling compared to those found in Gram-negative bacteria. The lack of the OM components together with the incapacity to detect associated extracellular pili raise questions about the assembly mode of a conducting channel that crosses the cell-wall and ensures a cell-to-cell contact. Several lines of evidence support the idea that Conj-T4SSs from Gram-positive bacteria emerged from diderm Conj-T4SSs and form a “minimized” system spanning the cytoplasmic membrane and the murein cell wall [8]. The best-characterized systems are those encoded by conjugative plasmids pCW3 from *Clostridium perfringens*, and pIP501 and pCF10 from *Enterococcus faecalis* [9–11]. Based on the structural and genetic information gathered from previous studies, it is assumed that the Conj-T4SSs from Gram-positive bacteria are essentially composed of subunits related to components recovered in Gram-negative T4SS. These ones comprise (i) VirB4 and VirD4-like energetic components, (ii) VirB3-, VirB6- and VirB8-like membrane subunits that assemble in the cytoplasmic membrane to form the membrane platform, (iii) VirB1-like transglycosylase and (iv) surface adhesins crucial in cell-to-cell contact or cells aggregation [12–17]. Finally, Conj-T4SSs from Gram-positive bacteria also comprise a set of accessory proteins with undefined functions [18]. However, the limited structural and mechanistic information on these components restrict our understanding of their mode of assembly and function.

Despite the similitudes in the early steps of DNA processing, fundamental differences exist in terms of channel architecture between Gram-negative and -positive Conj-T4SSs. Indeed, two components, a VirB1-like transglycosylase and a VirB8-like protein were predicted to extend their C-terminal domains along the exterior face of the cytoplasmic membrane and form a protruding channel for DNA translocation through the cell wall [8]. Unfortunately, such structural organization was not fully compatible with the crystal structures of the VirB8-like subunits TcpC and TraM from pCW3 and pIP501, respectively [19, 20]. Nevertheless, is it not excluded that channel establishment depends on VirB8-like interaction with other cytoplasmic membrane subunits including the VirB1-like transglycosylase.

Two types of mobile genetic elements are autonomous for their conjugative transfer: plasmids and ICEs. With the expansion of available genomes, it emerges that ICEs are the most widespread mobile genetic elements that use conjugation to propagate autonomously intra or interspecies [21, 22]. The 28 kb ICE*St3* from *Streptococcus thermophilus* is the prototype of the ICE*St3*/ICE*Bs1*/Tn*916* superfamily, the most widespread superfamily among Streptococci [23]. A large number of derivatives from ICE*St3*/ICE*Bs1*/Tn*916* contribute to the spread of various antimicrobial resistance genes in human or zoonotic pathogens including tetracycline and macrolide resistance genes [24, 25]. ICE*St3* was shown to be self-transferable intra- or inter-specifically including towards the human pathogen *Streptococcus pyogenes* and the opportunistic pathogen *E. faecalis* [26]. The ICE*St3* conjugation module encodes 14 proteins (OrfA to OrfN) [27]. Several of these proteins are homologous to the IMP components of the Gram-negative conjugative T4SS, whereas others (OrfB, E, F, H, L, M, N) are not. The presence of these latter Orfs with undefined structural homologues and functional information suggests distinct structural and assembly features of the conjugative apparatus of ICE*St3* and may reflect a specific functioning mechanism.

This study belongs to a multiapproach project that aims to understand the architecture and the assembly mode of Conj-T4SS from ICEs found in Gram-positive bacteria. Here, by using the ICE*St3* as biological model, we focus on the structure-function analysis of several proteins of the Conj-T4SS. Among them, OrfG shares a low sequence identity (22%) with TcpC, the VirB8-like component of pCW3. A structural analysis revealed that the OrfG central domain adopts a NTF2-like fold and likely presents more structural similarities with VirB8-like proteins TcpC and TraM from Gram-positive bacteria than Gram-negative VirB8 proteins. We used structural alignment tools to identify different structural classes among VirB8-like proteins. Our *in-silico* analysis revealed that Gram-positive VirB8-like proteins form a distinct class from Gram-negative ones, supporting a different evolutionary pathway. While already described soluble domains of Gram-positive VirB8-like proteins were observed as monomeric in solution, OrfG was found to self-assemble into 6-mer oligomers. The significance of this quaternary structure is discussed in view of the role of this protein in Gram positive Conj-T4SSs.

## Results

### OrfG belongs to the TcpC family

Previous studies showed that the integrative and conjugative element ICE*St3* from *S. thermophilus* is an autonomous element able to self-transfer by conjugation intra and interspecifically [26]. Bioinformatic analyses of ICE*St3* predicted the presence of a conjugation module that harbors 14 genes. Among them, *orfJ* was recently showed to encode the relaxase that belongs to the MobT superfamily [28]. In this study, we focused on the structural and biochemical studies of a putative transfer protein OrfG. Blastp analysis of OrfG indicated that this protein is composed of a TcpC domain (pfam12642) from position 98-316 (e-value 7.4e^−39^) (Fig. 1A). This domain is homologous to that of several TcpC conjugal transfer proteins encoded by diverse ICEs or conjugative plasmids, including the VirB8-like subunit TcpC from pCW3. The archetypal TcpC from pCW3 is a membrane protein that was shown to be essential for the conjugation of pCW3 from *C. perfringens* [20]. TcpC consists of an N-terminal cytoplasmic domain followed by a transmembrane domain (TMD) which encompasses residues 57-79 and a soluble domain extended in the cell-wall. Despite a poor sequence identity/similarity between OrfG and TcpC (23% identity/38% similarity) in their TcpC domain, *in silico* analysis using HHPRED and CCTOP revealed that OrfG shares the same domain organization as observed for TcpC. Thus, OrfG is composed of an N-terminal tail followed by a TMD (residues 37 to 59) and a soluble domain that we called OrfG_64-331_ (Fig. 1A).

**Figure 1.**
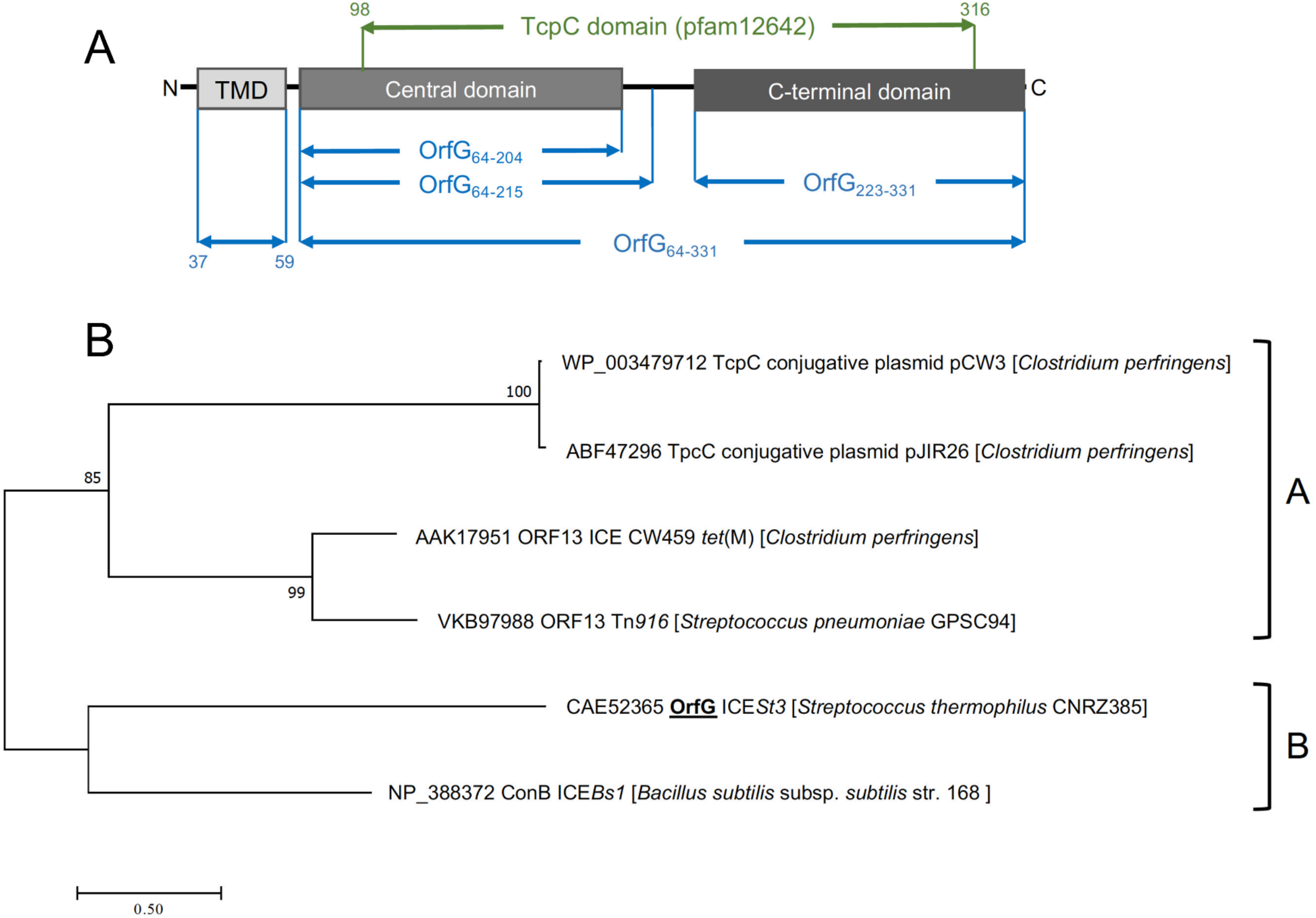
OrfG belongs to the TcpC superfamily. (A) Schematic representation of OrfG subdomains and their boundaries. (B) Phylogenetic tree of representative members from the TcpC superfamily including OrfG. This tree was inferred by using the Maximum Likelihood method based on the JTT matrix-based model [54]. The percentage of trees in which the associated taxa clustered together is shown next to the branches. The tree is drawn to scale, with branch lengths measured as the number of substitutions per site. All positions containing gaps and missing data were eliminated. There was a total of 284 positions in the final dataset. Evolutionary analyses were conducted using MEGA7 [51].

Our phylogenetic analysis of OrfG and TcpC orthologs indicated that OrfG is more closely related to the VirB8-like protein (conB) from ICE*Bs1* [29] than to the other TcpC proteins (including TcpC from pCW3) [20]. This is consistent with the existence of at least 2 sub-families of TcpC proteins (Fig. 1B). Given the similarities in domain organization between OrfG and TcpC, their distant evolutionary relationship motivated us to focus on the structural and biochemical characterization of OrfG.

### OrfG central domain (OrfG_64-204_) displays a VirB8-like structure

Owing to the lack of structural data on Gram-positive Conj-T4SS components especially those involved in ICE conjugation, we envisaged structure determination of OrfG by X-ray diffraction. For this purpose, we purified to homogeneity a large amount of the OrfG soluble domain (OrfG_64-331_: from Ser64 to Asp331). Despite an easy growing of crystals in multiple and different conditions, their poor X-ray diffraction power prevented the acquisition of data of sufficient quality. To overcome these technical issues, and based on the secondary structure prediction of OrfG as well as the sequence alignment between OrfG and TcpC, we chose to express and purify to homogeneity the separate central (OrfG_64-204_: from Ser64 to Ala204) and C-terminal (OrfG_223-331_: from Ala223 to Asp331) subdomains (Fig. 1A). Crystallization trials conducted on both forms resulted in suitable tetragonal crystals for OrfG_64-204_ while no positive result was obtained for OrfG_223-331_. Experimental phases were determined by using a single-wavelength anomalous dispersion method (SAD) on crystals soaked with an osmium derivative. A native crystal allowed final model refinement to 1.75 Å resolution (*R_free_* = 21.2 %). Detailed data and refinement statistics are given in Table S1.

The polypeptide chain adopts an alpha+beta fold, composed of two α-helices named α_1_ and α_2_, followed by a β-sheet of four strands (β_1_ to β_4_), the last of which has a marked disruption into two subparts β_4a_ and β_4b_ (Fig. 2). Cross-comparisons performed by DALI [30] and PDBefold [31] classify the spark OrfG central domain in the NTF2-like superfamily. Its folding pattern mainly consists in a highly curved antiparallel beta sheet that wraps around a central alpha helix surrounded by shorter additional ones. As expected from sequence analysis, the closest VirB8-like structural neighbours of OrfG_64-204_ is the TcpC central domain (TcpC_104-231_, i.e. the first domain of TcpC_99-309_, PDB entry 3ub1 [20]), with an rmsd of 1.77 Å for 92 aligned amino acids. This search also highlighted the structure of the C-terminal domain of TraM from pIP501 (TraM_190-322_, PDB entry 4ec6 [19], rmsd of 1.90 Å for 96 aligned amino acids) as a close structural homolog (Fig. 2). The folds of TcpC_104-231_, TraM_190-322_ and OrfG_64-204_ differ by the length of their secondary structures and connecting loops. Furthermore, TcpC_104-231_ has an additional helix α_3_ prior to the β-sheet. Both TcpC and TraM were described as Gram-positive equivalents of VirB8 from VirB/D4 T4SS from *A. tumefaciens*. Despite their low sequence identity and their different domain organization, it was proposed that proteins from Gram-positive Conj-T4SSs that adopt a NTF2-like fold form the “VirB8-like” family [23]. This family also includes the soluble domain of the conjugative component TraH from pIP501 (TraH_57-183_, PDB entry 5aiw [32]) (Fig. 2). Since the structure of OrfG_64-204_ belongs to the NTF2-like superfamily as well, we propose that OrfG from ICE*St3* is a new member of the VirB8-like family found in Gram-positive Conj-T4SS.

**Figure 2.**
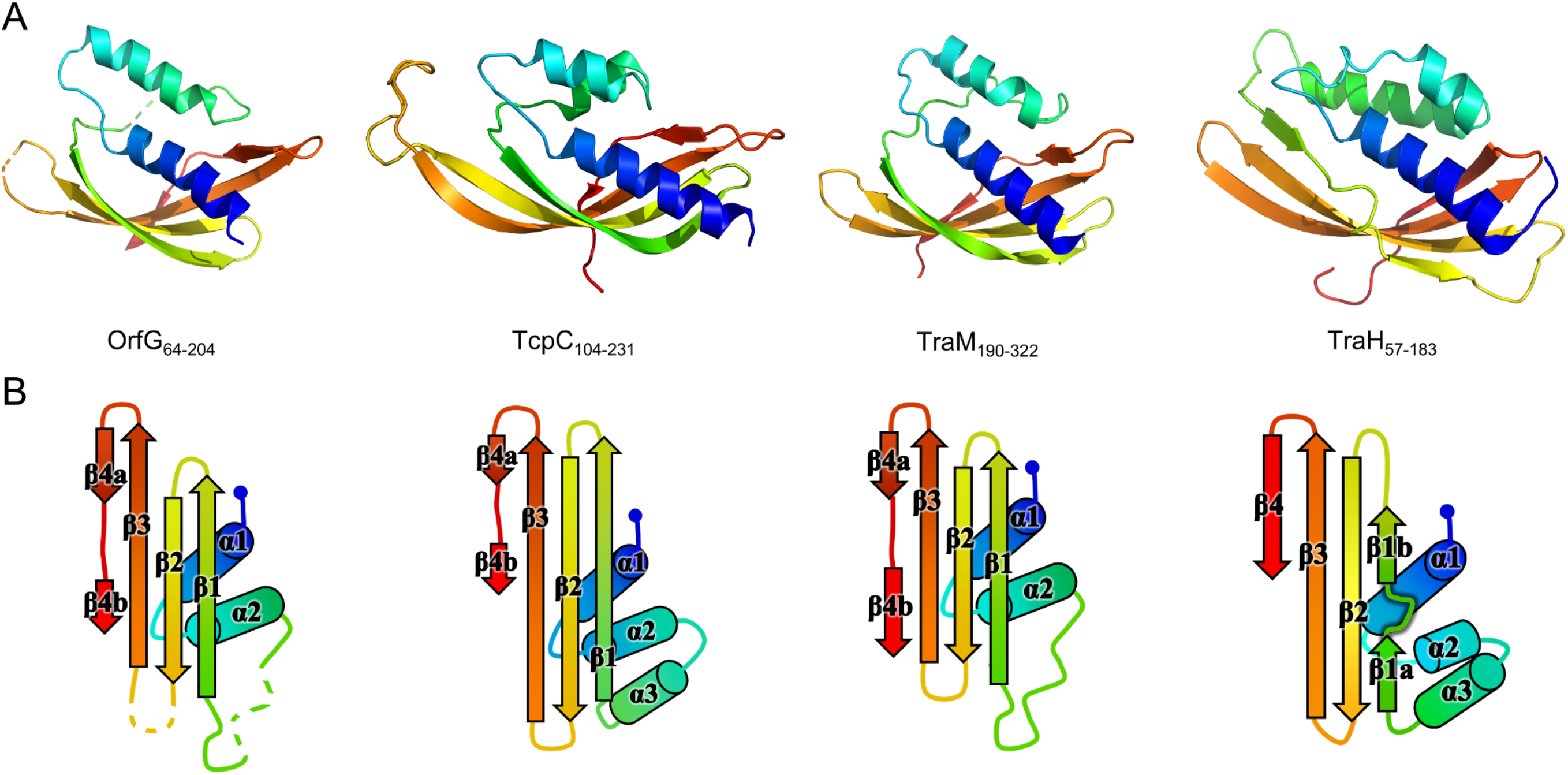
Structures of OrfG_64-204_, TcpC_104-231_, TraM_190-322_ and TraH_57-183_. From left to right respectively, the four monomers (PDB id 6zgn, 4ec6, 3ub1 and 5aiw) are displayed in the same orientation and represented as rainbow-colored cartoons, with the N-terminal end of the polypeptide chain in blue and the C-terminal end in red (A). Their topology is schematized with the same rainbow-color code and the name of the secondary structures (B).

### Structural comparison of available VirB8-like structures

Unexpectedly, known structures of VirB8-like proteins show subdomains that harbor an overall fold similar to the unique NTF2-like domain of Gram-negative VirB8s. This observation sparked our interest in a structural comparison of these proteins. A previous study using sequence analysis, secondary structure prediction and domain composition of a large set of sequences distinguished three classes of proteins [19]. Here we conducted a distinct analysis that focused on the available three-dimensional structures of the NTF2-like domain of these proteins, with the aim to point out the shared features and the specific characteristics of both Gram-positive and Gram-negative proteins. It was based on the superposition of their atomic coordinates, independently of their amino acid sequence. The results emphasized a surprising diversity despite the simplicity and small size of the NTF2-like fold.

The set of Gram-negative VirB8 structures encompasses 13 independent entries in the protein database (Protein Data Bank). That of known Gram-positive VirB8-like structures is poorer: the crystallographic analysis of OrfG_64-204_ brings their number to four. The NTF2-like pattern is globally conserved in the 17 structures. On average, their fold consists in three antiparallel alpha helices followed by an antiparallel four-stranded sheet, for which we propose the nomenclature α_1_α_2_α_3_β_1_β_2_β_3_β_4_. However, the length of the secondary structures (especially beta strands) and their relative orientation, the curvature of the beta sheet, plus possible decorations inserted in the minimal fold bring enough diversity to make the structural superimposition delicate. We found the mTM-align algorithm as the most suited one to deal with these differences. The global superimposition of the 17 structures and the corresponding structural alignment provided by mTM-align were analyzed the using an in-house developed script to allow their visualization in Pymol (see Materials and Methods) (Fig. 3). The structure-based tree obtained using mTM-align indicated that Gram-positive VirB8-like structures clearly separate from the group formed by the Gram-negative ones (Fig. 4). The tree also shows different classes in both Gram-negative and Gram-positive VirB8-like structures, which seem to have a real structural meaning since they contain members whose sequences share modest to low similarities (Table S2). At least three classes appear in the Gram-negative VirB8s (named I^−^, II^−^ and III^−^ for the convenience of their description) and one in the Gram-positive VirB8-like structures (I^+^), while the three other proteins and did not belong to identified classes (Fig. 4). In the Gram-negative group, the three classes differ by the way their helix α_3_ is disrupted in two parts α_3a_ and α_3b_, separated by a protruding loop of 4 residues with two distinct conformations in classes I^−^ and II^−^, or by a strong kink in class III^−^ (Fig. 4). This disruption of α_3_ is not a distinctive trait of the Gram-negative proteins since CagV from *Helicobacter pylori* (PDB entry 6iqt [33]), not assigned to a class, has a straight helix. All members of classes I^−^ and II^−^ have a small additional helix α_4_ between β_3_ and β_4_, which is involved in protein dimerization, while members of class III^−^ have a loop instead. Furthermore, class III^−^ and CagV display a bulge that separates β_1_ in two parts β_1a_ and β_1b_ (Fig. 4).

**Figure 3.**
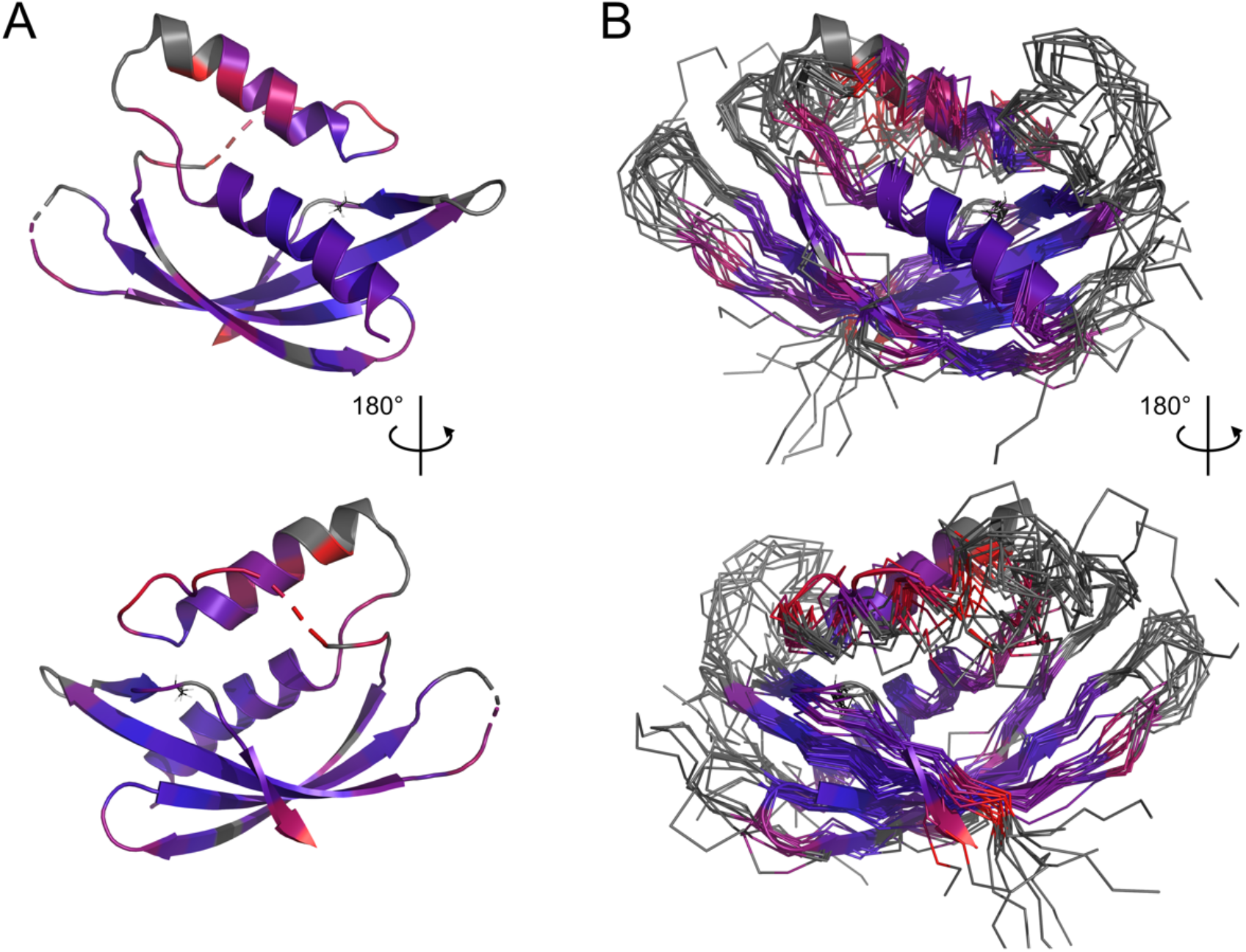
Comparison of 17 representative three-dimensional structures of VirB8 and VirB8-like proteins. The structure of OrfG_64-204_ (PDB id 6zgn) is represented as a cartoon (A), with its amino acids colored according to the quality of the superposition observed within a set of 17 selected structures. For each position, a gradient from violet to red is used depending on the low to high values of the root mean square calculated on the distance between all the possible pairs of the 17 superposed Cα carbons at this position (grey means that mTM-Align found at least one structure impossible to align). The amino acid shown with lines (Val191 in OrfG) and localised in β_4_ facing α_1_ is the only one whose side chain shares a similar nature in all structures, while there is no conserved residue. The whole set of structures (PDB ids 2cc3, 3ub1(limited to TcpC_104-231_), 3wz3, 3wz4, 4akz, 4ec6, 4jf8, 4kz1, 4lso, 4mei, 4nhf, 4o3v, 5aiw, 5cnl, 5i97, 6iqt, 6zgn) represented as a wire using the same color-code (B) reveal resemblances and differences in their NTF2-like folds.

**Figure 4.**
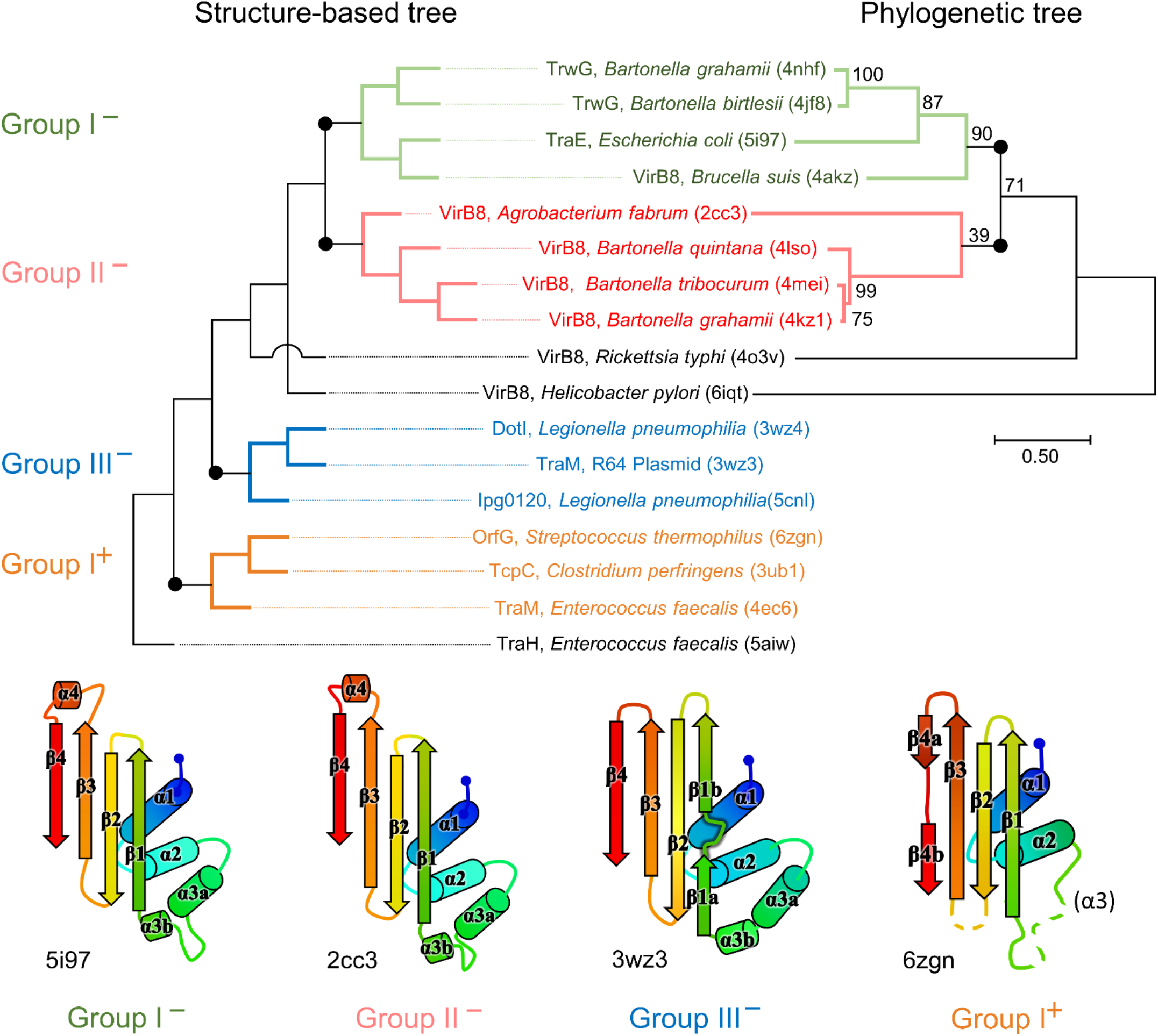
Structural and phylogenetic trees of Gram-negative VirB8 and Gram-positive VirB8-like proteins. Structure-based tree derived from the optimized superposition of the atomic coordinates (left). 17 representative structures were retrieved from the PDB and compared by mTM-align [52]. The name of the protein, its source and PDB id are indicated for each protein. Members of class I^−^ are in green, class II^−^ in red, class III^−^ in blue, and class I^+^ in orange where the (^−^) and (^+^) correspond to VirB8-like structures from Gram-negative and -positive bacteria, respectively. Secondary structure topology observed within each class is schematized at the bottom of the figure. Among the proteins compared in the structural analysis, all the sequences that belong to the PFAM group PF04335 were subjected to multiple sequence alignment and neighbor-joining tree building. The result is shown on the right-hand side of the figure. Numbers at nodes indicate the bootstrap values as percentage (1,000 replicates). Scale bar indicates the number of amino acid differences per site.

Strikingly, the four Gram-positive VirB8-like proteins have no specific structural feature that unambiguously distinguishes them from the Gram-negative ones. However, the overall structure superposition separated them in the structural tree, which seems to be explained by the small differences disseminated along their polypeptide chains (Fig. 4). Indeed, OrfG_64-204_ has no α_3_ (replaced by an unobservable region) and a strand β_4_ that clearly splits in two short portions β_4a_ and β_4b_ separated by a bulge that ends with Pro200 (Fig. 2). By comparison, TraM_190-322_ (PDB entry 4ec6) has a shorter α_2_, also misses α_3_ and has a slightly longer loop between β_1_ and β_2_. In contrast, TcpC_104-231_ (PDB entry 3ub1) possesses α_3_ and has much longer strands β_1_, β_2_ and β_3_, together with a longer loop between them. TraH_57-183_ is the most distant structure, as it is the only one with a continuous strand β_4_ with no bulge insertion, while on the contrary β_1_ possesses two parts β_1a_ and β_1b_ with a bulge in between (PDB entry 5aiw) (Fig. 2), more similar to some Gram-negative proteins (class III^−^ and CagV). Furthermore, TraH has a pronounced helix α_3_ with four turns and a long loop towards β_1_.

The most distinguishable difference between Gram-negative and Gram-positive VirB8 proteins lies in the length of the connection between β_3_ and β_4_. It has 10 to 17 amino acids in Gram-negative VirB8s (including α4 in classes I^−^ and II^−^), and only 4 to 7 in Gram-positive ones. However, deleting all these atoms in the 17 structure files prior to calculation of the structural tree resulted in the same classification, with only minor modifications. Thus, it seems that Gram-positive and -negative proteins separate according to details evenly distributed across the structures instead of localized marked traits that would characterize each group.

The script that we developed to analyse the structural alignment of the 17 structures easily identified the conserved or similar residues within the different sets of proteins that we considered. Interestingly, we found no conserved residue common to all 17 structures. In the subset of the 13 Gram-negative VirB8 structures, once again, no amino acid is conserved. On the contrary, up to 30 residues are conserved within each Gram-negative class, among which many form hydrophobic contacts where the β-sheet faces α_1_ (Fig. S1). By comparison, in the four Gram-positive VirB8-like structures, only six residues share similar natures and constitute a hydrophobic patch within the core of the protein near the C-terminal side of α_1_. In addition, it is worthy to notice the presence of a conserved glutamate residue (Glu154 in OrfG) at the end of β_1_, with its side chain towards the surface. Its conservation might be the result of happenstance due to the low number of analyzed structures and no apparent involvement in any observed assemblies. As for Gram-negative VirB8s, only a subset of Gram-positive VirB8-like proteins display more conserved residues (designated as Class I^+^) (Fig. S1). The absence of any signature was already noticed in the NTF2-like superfamily [34]. However, it is surprising that, despite probably having the same function, this observation also applies to the restricted subset of NTF2-like superfamily proteins that constitute the VirB8-like proteins.

### Crystal packing of VirB8-like proteins raises the question of their multimerization

In the tetragonal crystal, the OrfG_64-204_ monomer interacts with symmetry-related partners, among which one forms a questionable pair with a nice arrangement of the facing alpha helices (Fig. 5A). However, the interaction involves a low surface area of 1 600 Å^2^, or only 12% of the solvent accessible surface, as determined by PISA [35], which rejects the arrangement as an effective quaternary structure. As well, we struggled to find an explanation to the resulting location of both N-termini in this association. Indeed, it would place the transmembrane domain of each OrfG monomer at opposite sides of the dimer, but this argument should be used with caution since 27 N-terminal residues are missing in the model with respect to the crystallized protein. Altogether, the pair of OrfG_64-204_ monomers observed in the crystal is not a convincing proof of the the existence of a dimer with a biological relevance.

**Figure 5.**
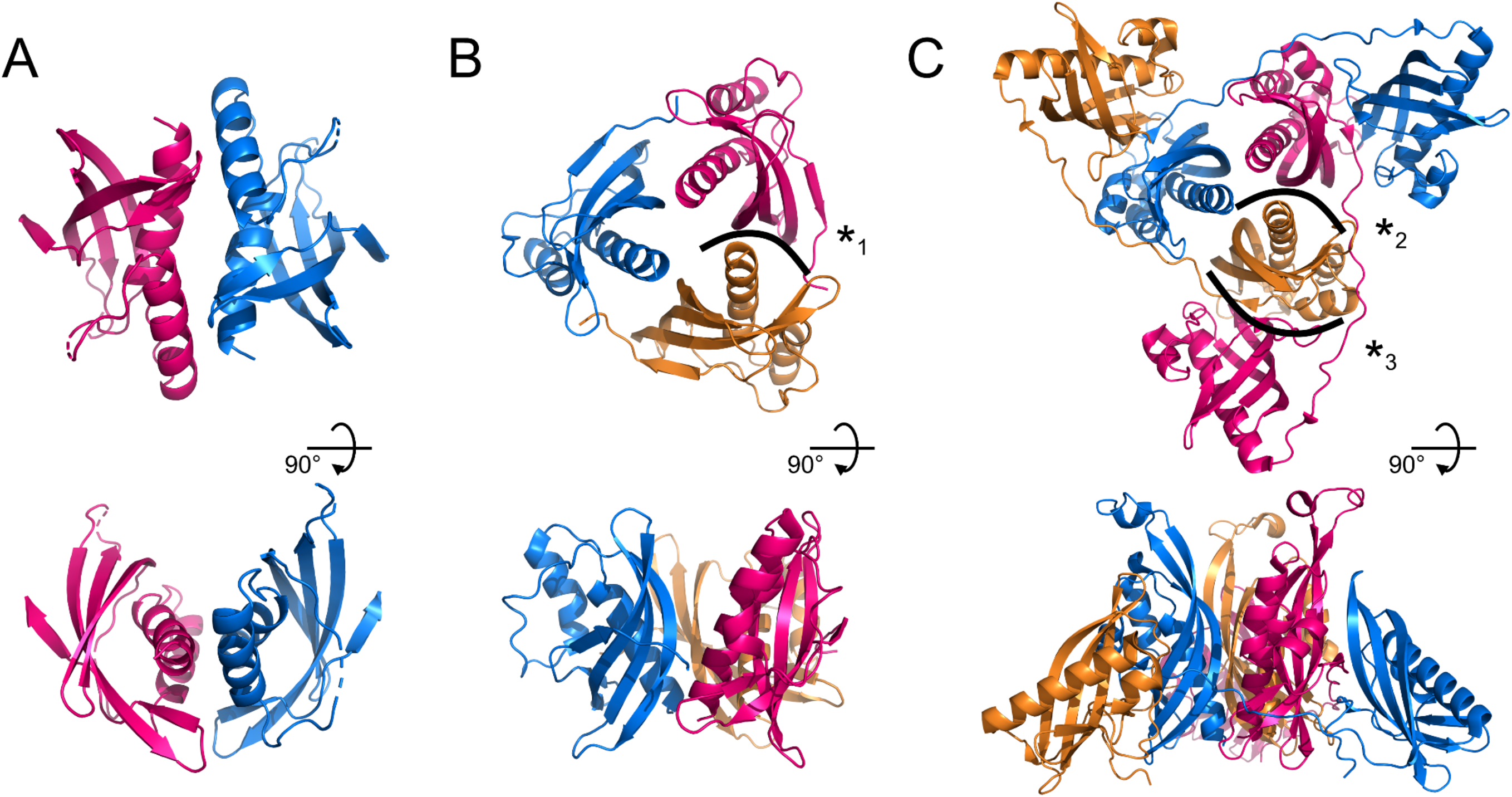
Assemblies of Gram-positive VirB8-like proteins observed in the crystals. A crystallographic 2-fold axis relates two monomers of OrfG_64-204_ (PDB id 6zgn) (A), while TraM_190-322_ (4ec6) (B) and TcpC_99-309_ (3ub1) (C) form trimers. Each assembly is shown perpendicular (top) and parallel (bottom) to its symmetry axis. The interface with the highest surface area, here represented for OrfG_64-204_, is resolutely rejected by PISA [35]. The interface (*1) in TraM_190-322_ extends on ~300 Å^2^ and involves 2 to 9 hydrogen bonds or salt bridges depending on the observed monomer (as calculated by PISA for residues 214 to 318 of 4ec6). The equivalent interface in TcpC_99-309_ (*2) represents ~430 Å^2^ (PISA on residues 104 to 228 of 3ub1) and 6 to 9 hydrogen bonds or salt bridge depending on the monomer. It is by far slighter than the interface (*3) observed between the central domain (TcpC_104-231_) of one monomer and the C-terminal domain (TcpC_239–354_) of a second one, which extends on ~980 Å^2^ and involves 13 to 14 hydrogen bonds or salt bridges depending on the observed monomer. The structure-based sequence alignment of OrfG, TcpC and TraM available in Figure 6 shows residues involved in these interactions.

Analysis of structural similarity positioned OrfG_64-204_ in the same class as TraM_190-322_ and TcpC_104-231_, while TraH_57-183_ is more distant (Fig. 4). The proximity of these three proteins raises once again the question of a possible multimerization of OrfG. Indeed, both TraM_190-322_ and TcpC_99-309_ form trimers in their crystal assembly (Fig. 5B and C). Yet, TraM_190-322_ corresponds to a single domain, like the fragment of OrfG that has been crystallized (OrfG_64-204_), while TcpC_99-309_ has two subdomains equivalent to the central and C-terminal domains of OrfG (OrfG_64-331_, no structure available). Do the protein interfaces share homology that allows prediction of an equivalent assembly of OrfG? Alternatively, does the latter has specificities that would result in a different oligomeric form? Comparison of the trimers formed by TraM_190-322_ and TcpC_99-309_ emphasizes a surprising similarity in the positioning of their common domain (Fig. 5B and C). TraM_190-322_ and the central domain of TcpC (TcpC_104-231_) bring their N-termini near the 3-fold axis that defines the trimer. Interactions that build this assembly are mainly located at the α_1_ N-terminal moiety of one partner and near the β_1_ C-terminal end of the facing partner, in a circular pattern of interactions. Detailed analyses are provided in the papers that describe these crystallographic structures [19, 20]. Briefly, TraM_190-322_ hides 14% of the calculated solvent-accessible surface area of one isolated monomer in this contact, which involves three hydrogen bonds and several van der Waals contacts (Fig. 5B). Consistently with this observation, the weakness of this interaction gives little credibility to the existence of a trimer in solution. However, its existence in the crystal makes sense with respect to the prediction of a coiled-coil motif formed by some thirty residues that precede the NTF2-like domain, which could stabilize a trimeric assembly of TraM [19]. In the case of TcpC_99-309_, the equivalent interface between the central domains TcpC_104-231_ is a bit wider and based on more hydrogen bonds (a total of eight) [20] (Fig. 5C). The striking observation that arises from a precise comparison of these interfaces is their total lack of commonality, except for the region they concern. Two particular examples illustrate this conclusion. First, in TraM_190-322_, the side chain of Lys217 at the beginning of α_1_ interacts with Tyr269 in β_1_ of the partner, via van der Waals contact with its aromatic ring and the addition of a hydrogen bond with its main chain on four of the six instances observed in the asymmetric unit. At the equivalent positions of TcpC_99-309_ (Fig. 6), Gln108 and especially Phe161 have different orientations so that instead Gln108 interacts with Glu106 at the beginning of α_1_ in the neighbouring monomer. The second example concerns Tyr224 (α_1_) in TraM_190-322_, the side chain of which forms a hydrogen bond with Asn285 (β_2_) of the associated monomer. In TcpC_99-309_, the residue Glu115 at the equivalent position in α_1_ prefers an interaction with Ser159 in β_1_ of the facing monomer. As a conclusion, the driving force that governs the trimer formation does not originate from conserved residues at this interface. However, the observation of similar multimeric forms with similar arrangements is suggestive of their probable relevance. Structure and sequence comparisons with OrfG_64-204_ revealed no conservation of interacting pairs of residues (neither with TcpC_99-309_ nor with TraM_190-322_) but they also revealed no element that would prevent formation of the same interaction (Fig. 6).

**Figure 6.**
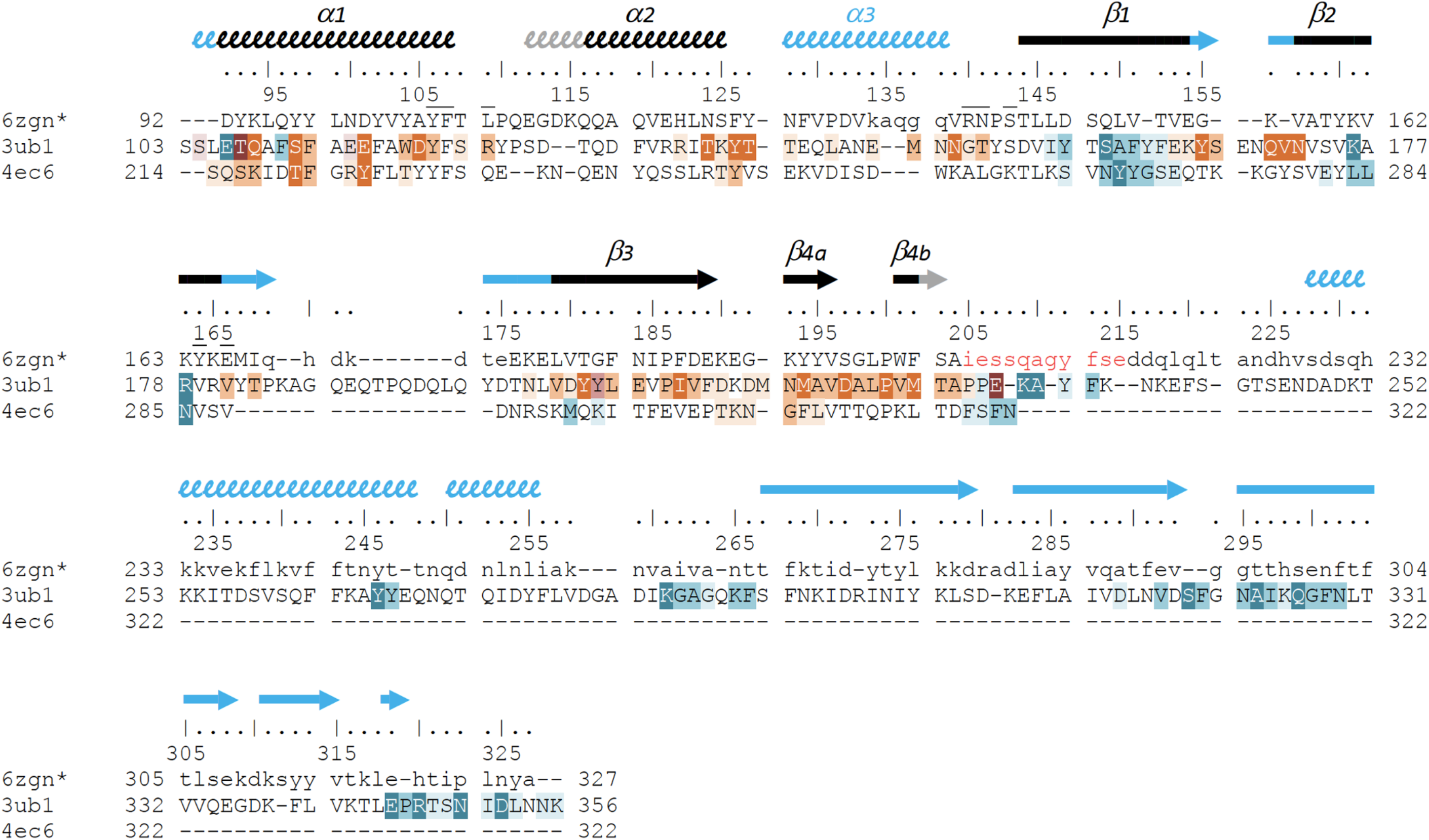
Structure-based sequence alignment of three Gram-positive VirB8-like proteins. The sequence alignment was generated by mTM-align and manually modified, based on the three structures of OrfG_64-204_ (pdb entry 6zgn), TcpC_99-359_ (3ub1) and TraM_190-322_ (4ec6). Residues of OrfG_64-204_ that were not observed in the electron density are shown as lowercases, as well as residues 205-327 equivalent to the C-terminal domain of TcpC and for which no structure is known yet. Numbering above the sequences corresponds to OrfG. Secondary structures are represented by arrows (β-strands) and squiggles (α-helices), in black when shared by OrfG and TcpC, in grey when only present in OrfG, and in cyan when only present in TcpC. Residues of TraM and TcpC involved in the trimer assemblies are highlighted, in orange in one partner and in blue in the facing monomer: white letters with dark background are for residues involved in hydrogen bonds in all interface instances in the asymmetric units, while dark letters with lighter background are involved in contact in at least one interface (medium background when at least 50% of the residue is buried in the contact, light background if less than 50% is buried). Residues of OrfG_64-204_ that delineate the pocket in which an unknown ligand is bound (see Fig. S2) are marked with an overbar.

No trimer is observed in the crystal form of OrfG_64-204_. Deletion of the C-terminal domain is a plausible explanation but it is not appropriate for TraM_190-322_. A closer analysis of the different structures shows that TraM_190-322_ has a C-terminal extension of five residues after β_4b_, which follows the same direction as the first residues of the linker between the two domains of TcpC_99-309_. Its penultimate residue Phe321 positions its side chain in a hydrophobic environment formed by several aromatic residues in α_1_, α_2_ and β_4a_ of the facing monomer. In TcpC_99-309_, the linker residue Lys233 also contributes to stabilization of the trimer, but via a hydrogen bond with the polypeptide main chain following α_2_. In OrfG, the equivalent Gln209 could play the same role (Fig. 6). In order to evaluate the contribution of this extension, we solved the structure of OrfG_64-215_. However, no change was induced, neither in the crystallization conditions nor in monomer packing in the crystal (data not shown) demonstrating that the linker was not sufficient to promote OrfG_64-215_ multimerization.

The crystal structure of TcpC soluble domain (TcpC_99-309_) shows that its C-terminal domain (TcpC_239–354_) also belongs to the NTF2-like superfamily [20]. It has a peripheral position in the trimeric assembly, far from the N-terminal domain it is linked with (TcpC_104-231_). It gives a monomer shaped like a telephone receiver, three of them being interlaced to constitute the trimer (Fig. 5C). This organization mode results in a large interface (~1000 Å^2^) between the central domain of one monomer and the C-terminal domain of the partner. Thirteen hydrogen bonds are formed, together with hydrophobic interactions. It alone accounts for much more than the interactions formed between the central domains and described above. However, this convincing arrangement, which buries 24% of the accessible surface area of each monomer, was not observed in solution [20]. The authors proposed that the deleted N-terminal residues, in particular a transmembrane domain, could explain this discrepancy. The similarity between OrfG and TcpC in their domain organization and in the structure of their central domain, despite a low sequence identity, suggests a similar overall structure of the whole protein, and a similar oligomeric state. The missing linker and C-terminal domain probably hinder trimerization in the crystallization conditions used.

### In vitro self-oligomerization of OrfG_64-331_ and its subdomains

The closest structural homologs to OrfG_64-204_ form trimers in their crystal arrangement (Fig. 5B and C). Thus, it was proposed that trimers constitute the functional biological assembly of both proteins [19, 20]. These observations strongly call for the assessment of the oligomeric state of OrfG_64-204_ in solution. Protein multimerization can be monitored by chemical cross-linking. We performed *in vitro* chemical cross-linking experiments with a constant amount of the protein in presence of an increasing concentration of the cross-linking agent Paraformaldehyde (PFA). As observed in Figure 7A, in the presence of a low amount of PFA (0.05 to 1%), OrfG_64-331_ predominantly forms dimers. High molecular-mass oligomers of OrfG_64-331_ are visible at higher PFA concentrations (2 and 5%) demonstrating the predisposition of OrfG_64-331_ to oligomerize in solution. The ability of OrfG_64-331_ to self-assemble was confirmed by the analysis of the size exclusion chromatography (SEC) profile where the 31 kDa OrfG_64-331_ was eluted into two defined peaks. SDS-PAGE analysis showed that both peaks are exclusively composed by OrfG_64-331_ protein (Fig. 7B). These results suggest that OrfG_64-331_ adopts two oligomeric states in solution. To further examine the apparent existence of these two oligomeric states of OrfG_64-331_, we analyzed purified OrfG_64-331_ by SEC coupled to multiple angle light scattering (SEC-MALS) (Fig. 8A). The predominant form (97.3% of the total mass fraction) was eluted at 29 min and corresponds to the monomeric form of OrfG_64-331_ with a calculated MW of 32 kDa (theoretical MW=31 kDa). The less present form (2.7% of the total mass fraction) was eluted earlier at 25 min with a calculated MW of 172 ± 11 kDa which can be assigned as 6-mer of OrfG_64-331_ (186 kDa theoretical MW). SEC-MALS data confirm the ability of OrfG_64-331_ to form ordered homo-multimers in solution. Intriguingly, such behavior has not been reported for other characterized members of the Gram-positive VirB8-like proteins. In fact, AUC and SEC analysis performed for TcpC and TraH truncated for their TMD, respectively, revealed that both proteins were strictly monomeric in solution [20, 32]. TraM_190-322_ (TraM lacking its N-terminal domain and the TMD) was able to form oligomers only in the presence of cross-linking agents whereas DLS and SAXS analysis indicated a monomeric form of TraM in solution [19]. The knowledge that OrfG can assemble into ordered oligomers in solution led us to question whether OrfG central domain or OrfG C-terminal domain or both would promote/contribute to OrfG_64-331_ oligomerization. For that purpose, we investigate separately their capacity to self-assemble by using chemical cross-linking. Regarding OrfG_64-204_, the *in vitro* chemical cross-linking using PFA gave mostly a dimeric form when using at least 0.05% of PFA (Fig. 7C). At higher PFA concentrations, a fussy additional band with a higher mass was observed that might correspond to a trimeric form. On the contrary to OrfG_64-204_, the chemical cross-linking of OrfG_223-331_ leads to the formation of multiple oligomers of OrfG_223-331_ easily distinguished when analyzed by SDS-PAGE (Fig. 7E). These results suggest that both subdomains were independently able to form oligomers in solution. Interestingly, during SEC analysis of both OrfG_64-204_ and OrfG_223-331_, we observed the presence of two well-defined peaks as seen on the SEC profile of OrfG_64-331_ (Fig. 7B, D, F). These findings prompted us to characterize further the oligomerization of OrfG_64-204_ and OrfG_223-331_ using SEC-MALS. As observed in Figures 8B, C, both OrfG_64-204_ and OrfG_223-331_ form two oligomeric states. A predominant monomeric form was eluted at 36 min for OrfG_64-204_ and at 34 min for OrfG_223-331_ with a calculated molar mass of 17 kDa and 13 kDa, respectively. Additional early peaks were barely detected at 32.5 min and 31 min with calculated molar mass of 108± 8 kDa and 95 ± 9 kDa for OrfG_64-204_ and OrfG_223-331_, respectively. Interestingly, calculated molar masses for the ordered oligomers detected for the OrfG_64-331_ subdomains were compatible with 6-mer assembly for each subdomain (102 kDa and 78 kDa theoretical MW for OrfG_64-204_ and OrfG_223-331_, respectively).

**Figure 7.**
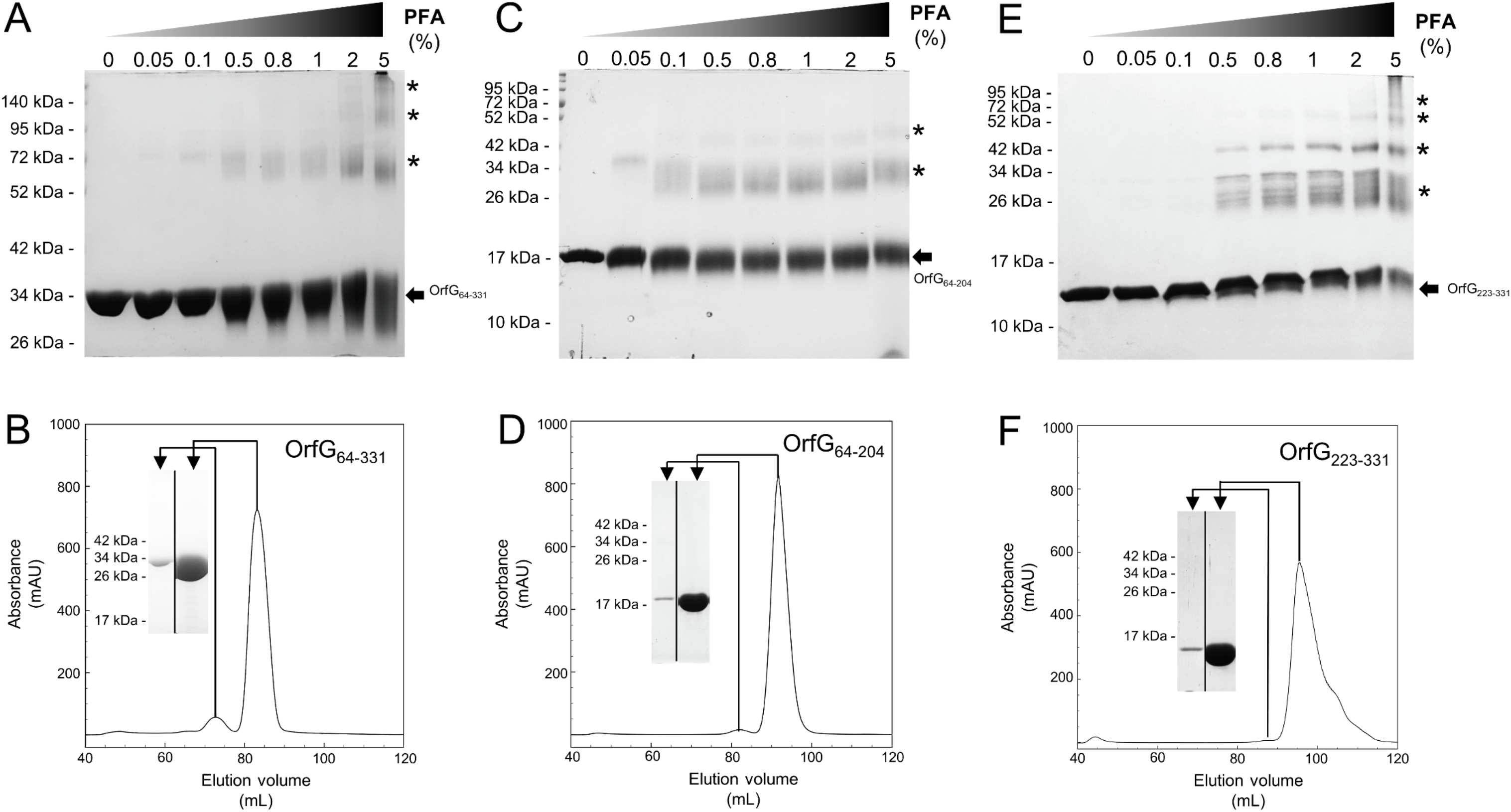
Analysis of oligomerization of native and truncated versions of OrfG_64-331_ by *in vitro* chemical cross-linking and size exclusion chromatography. *In vitro* chemical cross-linking of OrfG_64-331_ (A), OrfG_64-204_ (C) and OrfG_223-331_ (E). SDS-PAGE analysis of 3 μM of the purified OrfG_64-331_ and its truncated versions in absence or in presence of an increasing concentration (0.05 to 5%) of paraformaldehyde (PFA). Samples were loaded without heating treatment. The concentration of PFA used for each reaction is mentioned at the top of each column. Protein bands corresponding to each protein are indicated by black arrows. Asterisks indicate the identified oligomers stabilized by PFA cross-linking. Size exclusion chromatography (SEC) of the purified OrfG_64-331_ (B), OrfG_64-204_ (D) and OrfG_223-331_ (F) using Superdex 200 16/600. The elution volume (mL) is plotted on the x-axis and the 280-nm absorbance is plotted on the y-axis. An inset next to the elution profile corresponds to the analysis by SDS-PAGE of a unique fraction from each peak eluted during SEC purification. Electrophoretic separation shows that all identified peaks contain exclusively the analyzed protein. Molecular weight markers (in kDa) are indicated on the left of each SDS-PAGE pattern.

**Figure 8.**
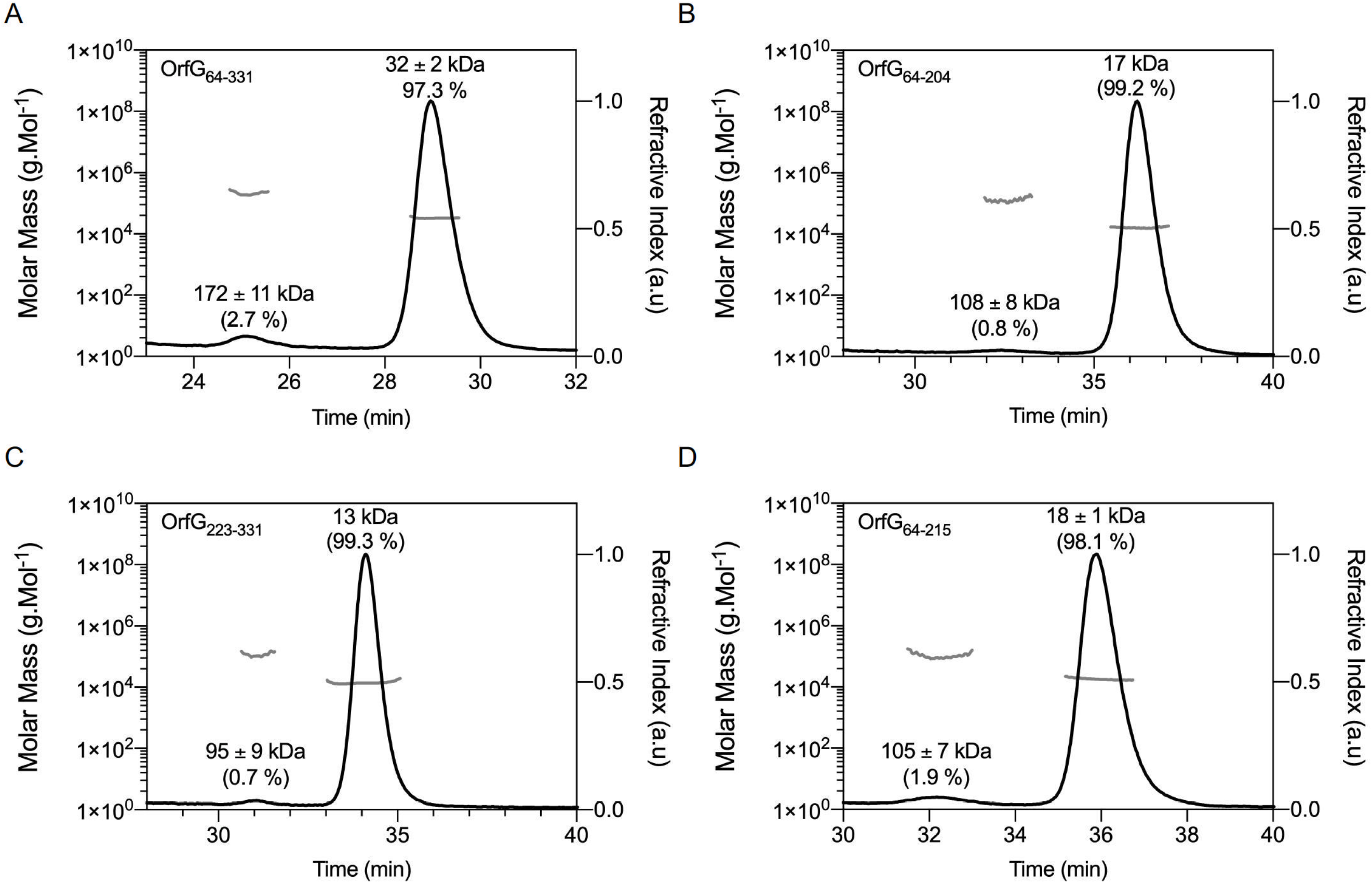
SEC-MALS analysis. Elution profile (black lines) of OrfG_64-331_ (A), OrfG_64-204_ (B), OrfG_223-331_ (C) and OrfG_64-215_ (D) are shown with the molecular weight calculated by MALS (gray lines). The elution time (min) is plotted on the x-axis. The molar mass (in logarithmic scale) is plotted in the first y-axis and the refractive index is plotted in the second y-axis. The molecular weight (in kDa) and the contribution in mass fraction (in %) of each visible peak are shown at the top of the corresponding peak.

## Discussion

During the last decades, a great effort was made to understand the mode of action of conjugative type IV secretion systems (Conj-T4SSs) due to their major role in antibiotic resistance spreading among bacteria. More attention was devoted to the study of Conj-T4SS from Gram-negative bacteria. Multiple structural information have been gained for individual components/proteins and isolated complexes improving our understanding of their architecture and assembly mode (reviewed in [5]). On the other side, Conj-T4SS from Gram-positive bacteria have been less investigated resulting in little knowledge on their architecture and on the molecular mechanisms underlying DNA transfer in these bacteria (reviewed in [9]).

In order to gain further information on the architecture and the assembly mode of Conj-T4SS from Gram-positive bacteria, we focused on the structural, biochemical and biophysical study of OrfG, a putative transfer protein encoded within the conjugative module of ICE*St3* from *S. thermophilus*. By using X-ray crystallography, we solved the structure of the central domain of OrfG. Structural analysis revealed that OrfG central domain adopts an NTF2-like fold, common to all known VirB8-like proteins in both Gram-negative and -positive Conj-T4SS. VirB8-like proteins are essential structural and functional component of Conj-T4SSs. Consequently, we propose that OrfG operates as a VirB8-like protein in the conjugative apparatus of ICE*St3*. The structure of OrfG central domain represents the first structure of a VirB8-like protein from ICEs described so far and rises to four the VirB8-like protein solved structures for Gram-positive Conj-T4SS. Notwithstanding the tricky task to identify VirB8-like proteins in Gram-positive Conj-T4SS clusters due to their low sequence identity, the growing number of VirB8-like structures indicates that it is a conserved protein in conjugation machines and underlines the similarity between protein complexes that govern the transport of mobile genetic elements across the cytoplasmic membrane in both Gram-negative and -positive bacteria.

OrfG central domain shares strong resemblance with the TcpC central domain of pCW3 from *C. perfringens*. Despite their low sequence similarity/identity, *in silico* analysis suggests that both OrfG and TcpC adopt the same domain organization with an N-terminal transmembrane domain (TMD) followed by two soluble subdomains. This domain organization appears to be conserved in a large number of putative transfer proteins from both conjugative plasmids and ICEs alike TcpC from pJIR26, Orf13 from ICECW459, Tn*916* and Tn*5397* and the VirB8-like protein (conB) from ICE*Bs1*. Our phylogenetic analysis comforts this observation showing that these proteins belong to the TcpC family (Fig. 1B).

In addition to TcpC and OrfG, two other transfer proteins, TraM and TraH from the conjugative plasmid pIP501 from *E. faecalis,* hold a NTF2-like domain in their C-terminal domains and were proposed to be VirB8-like proteins. Interestingly, both TraM and TraH adopt a different domain organization compared to TcpC and OrfG. In fact, TraM is composed of two soluble subdomains separated by a TMD while TraH is composed of an N-terminal TMD followed by a unique C-terminal subdomain. The variability in domain organization is restricted to VirB8-like proteins from Gram-positive bacteria since VirB8-like proteins from Gram-negative bacteria share a common domain organization with an N-terminal TMD followed by a single NTF2-like C-terminal periplasmic domain. The modularity/flexibility in domain organization perceived for Gram-positive VirB8-like proteins could be explained in part by the evolutionary scenario of conjugative systems. In fact, it was proposed that Gram-positive conjugative systems emerge from Gram-negative ones by gene deletion and such transfer was followed by diverse adaptive routes [36]. This significant variation could be originated from the varied cell envelope composition and thickness and/or the functional adaptation of conjugative systems regarding the nature of the donor and recipient cells as well as the variation of the conjugation environment [4]. Actually, most of the NTF2-like domains of the VirB8-like proteins from Gram-positive Conj-T4SS are localized in the cell-wall, such localization makes these domains more exposed to the environmental stress and this may contribute to their modularity compared to the periplasmic localization of VirB8-like protein in Gram-negative bacteria [37, 38].

Strikingly, although they are subtle, the differences observed when comparing the structures of all the NTF2-like domains of VirB8-like proteins lead again to distinguish Gram-positive and -negative bacteria. The tree based on structural similarities seems significant as it groups structures in a way comparable to the one obtained through phylogenetic analysis (when applicable i.e. for sequences that belong to the PFAM group PF04335, see Fig. 4). Moreover, this tree, strictly based on the spatial superposition of atoms without valuing sequence conservation, associates in a same class, proteins that share low sequence similarity. For instance, in the case of Gram-negative bacteria with more known structures, VirB8s from *A. tumefaciens* and from *Bartonella quintana* gather in class II^−^ although they only have 22% of sequence identity; Class III^−^ contains TraM from Plasmid R64 [39] and the IcmL-like protein from *Legionella pneumophila* despite only 13% of identity (Table S2). Thus, the partition of VirB8-like structures from Gram-negative bacteria on one side, and Gram-positive ones on the other side in the tree is highly significant since the low sequence similarity of the latter with any other could have distribute them anywhere in the tree. However, if clear in the tree, this distinction is less apparent when looking at the structures themselves, so that it is hard to define peculiar features that cause this distinction. Enrichment of the database by new structures of VirB8-like proteins from Gram-positive bacteria will probably bring elements that underpin this partition, as well as it will certainly draw several new classes in this part of the tree. The current distribution separates the monomeric *E. faecalis* TraH/Orf8 from the class I^+^ which contains TcpC from *C. perfringens* and TraM from *E. faecalis*. The presence of OrfG with these two members that share the same trimeric crystal packing questions the physiological assembly of this protein.

One of the most puzzling questions on the study of bacterial secretion systems including the Conj-T4SS concerns the oligomeric state of their components. In Conj-T4SS, VirB8 proteins are bitopic proteins of the cytoplasmic membrane. In Gram-negative bacteria, many studies suggested/supported that VirB8 function as a multimers. Most of the periplasmic domains of all studied VirB8 proteins adopt a dimeric assembly in solution and it is believed that these dimers are the forming subunits of higher-order multimers. Structural and biochemical data proposed that the full-length VirB8 protein self-assembles into a homo-multimers with C3 symmetry. Indeed, the stoichiometry of VirB8 was estimated to 12 in Conj-T4SS from *E. coli* R388 [40]. In line with, the purification of full-length TraE and TraM, a VirB8 homolog from pKM101 and R64, respectively, revealed that TraE and TraM forms hexamers in solution [39, 41]. Dissimilar to what was described in Gram-negative bacteria, VirB8-like proteins in Gram-positive bacteria are proposed to function as trimers. Up to date, no higher-order oligomers were obtained for VirB8-like proteins from Gram-positive bacteria supporting a different assembly mode compared to VirB8-like proteins recovered in Gram-negative bacteria. Nevertheless, this variation in the assembly mode do not amend the capacity of VirB8-like proteins to self-assemble following a C3 symmetry independently from the origin of the conjugative system. For Gram-positive VirB8-like proteins, TMD was proposed to trigger trimeric assembly since isolated TMD-less VirB8-like sub-domains were monomeric in solution. In addition, a predicted coiled-coil motif preceding the NTF2-like domain, not conserved in all VirB8-like proteins, seems to play an important role in the stabilization of the trimeric assembly [19]. In this context, our in-solution characterization revealed intriguing properties concerning OrfG: (i) significant but small fraction of OrfG soluble domain multimerize independently of its TMD and/or a coiled-coil domain since *in silico* analyses failed to found a predicted coiled-coil motif in the purified OrfG soluble domain, (ii) OrfG soluble domain has the ability to spontaneously assembles into 6-mer. This ordered oligomeric state observed in solution appears to be a physiological property of OrfG since SEC-MALS analysis performed on OrfG central and C-terminal sub-domains revealed that both form the same ordered-oligomers in solution. The low proportion of OrfG oligomers observed in solution could be explained by the absence of the TMD known to be essential for VirB8 oligomers stabilization [39, 41]. These data exposed a discrepancy on the number of the forming subunits between OrfG 6-mer oligomers and TcpC and TraM trimers. This could be explained by the fact that OrfG 6-mer oligomers could result from the dimerization of trimeric complexes. Another probable explanation lies on the ability of OrfG to adopt a different assembly mode not yet described for Gram-positive VirB8-like proteins.

Nevertheless, our results combined with published data obtained for TcpC, TraM and TraH consolidate the fact that VirB8-like proteins from Gram-positive bacteria act as multimeric entity on their biological systems but interrogates on their physiological stoichiometry that may depends on their dedicated systems.

VirB8 proteins from Gram-negative bacteria were considered to be interesting targets to develop specific inhibitors of Conj-T4SS by abolishing VirB8 dimerization. These inhibitors interact with a specific site localized in a structurally conserved groove not directly involved in the dimerization of VirB8 proteins [42, 43]. Since the inhibitor-binding interface is on the opposite site of dimerization interface, it was proposed that conformational changes on this groove directly affect VirB8 dimerization. Surprisingly, the electron density of OrfG central domain displayed an unassigned blob (Fig. S2) nicely docked in a pocket localized in a hydrophobic cavity that shares intriguing similarity with the groove identified in Gram-negative VirB8 proteins. In fact, residues located on the C-terminal end of α_1_ and β_4_ delimit both of the identified pockets. Thus, the localization of this groove suggests a probable oligomerization inhibition site for OrfG. The presence of this pocket in VirB8-like proteins could be explained by the nature of the NTF2-like fold. In fact, this fold is widely distributed in bacteria and is associated with various functions including enzymatically active and non-enzymatically active proteins. When extracellular, these proteins often possess non-catalytic ligand-binding activities [34].

## Materials and Methods

### Bacterial strains, media and chemicals

*Escherichia coli* DH5α was used for cloning procedures, BL21 (DE3) was used for protein expression and production. The genotype of each strain is listed in Table S3. Lysogenic Broth (LB) was used for bacterial culture. Kanamycin (50 μg/mL) was added to the culture medium.

### Plasmid construction

Plasmids used in this study are listed in Table S3. Polymerase Chain Reactions (PCR) were performed using a BioRad T100 Thermocycler thermal Cycler using Phusion High-fidelity DNA Polymerase (Thermo Scientific). *Streptococcus thermophilus* LMG18311 harboring ICE*St3* genomic DNA was used as template for PCR amplifications. Custom oligonucleotides (Eurogentec) used for cloning procedures are listed in Table S3. For protein expression and production, the DNA sequence encoding OrfG_64-331_, OrfG_64-204_, OrfG_64-_ 215 and OrfG_223-331_ were cloned into the pET28-TRX leading to pET28-TRX-OrfG_64-204_, pET28-TRX-OrfG_64-215_ and pET28-TRX-OrfG_223-331_, respectively. For *orfG_64-331_* and *orfG_223-331_*, restriction-ligation procedure was used for genes cloning. pET28-TRX-OrfG_64-204_ and pET28-TRX-OrfG_64-215_, plasmid derivatives were obtained by insertion of Stop codon at position 205 and 216 respectively using pET28-TRX-OrfG_64-204_ as template.

### Production and purification of OrfG_64-331_, OrfG_64-204_, OrfG_64-215_ and OrfG_223-331_

*E. coli* BL21 (DE3) cells carrying the pET28-TRX derivatives were grown on LB supplemented with Kanamycin (50 μg/mL), at 37°C to an OD_600_ ~ 0.5. Expression of the constructions was then induced by addition of IPTG (0,5mM) and cultures were pursued for 18 hours at 25°C. Cells were harvested, resuspended in lysis buffer (Tris-HCl 50 mM pH 8.0, NaCl 300 mM, EDTA 1 mM, lysozyme 50 μg/mL, phenylmethylsulfonyl fluoride (PMSF) 0,1 mM and submitted to 5 cycles of sonication. After addition of DNase (20 μg/mL) and MgCl_2_ (20 mM), the soluble fraction was obtained by centrifugation for 40 min at 16,000 × *g*. Recombinant proteins were purified by ion metal affinity chromatography using a 5-ml Nickel HisTrap™ HP Column on an ÄKTA prime apparatus (GE healthcare) pre-equilibrated in Tris-HCl 50 mM pH 8.0, NaCl 300 mM, Imidazole 10 mM (buffer A). After several washes in buffer A, 6×His tagged proteins were eluted in buffer A supplemented with 250 mM Imidazole, cleaved with the TEV protease (1 mg/mL) and dialyzed against Tris 50 mM, NaCl 300 mM pH 8.0 for 18 hours at 4°C and loaded onto a HisTrap™ HP pre-equilibrated in Buffer A, which selectively retains the TEV protease, the uncleaved, 6×His-tagged fusion proteins and contaminants. The native proteins were collected in the flow-through, concentrated on Centricons (Millipore; cutoff of 10 kDa), and passed through a Sephadex 200 26/60 column pre-equilibrated with Tris-HCl 50 mM pH 8.0, NaCl 100 mM or Hepes 20 mM, NaCl 100 mM.

### Crystallization and structure determination

Crystallization was conducted using the sitting drop, vapor diffusion method at 293 K. OrfG_64-204_ crystals were obtained by mixing 0.3 μL of protein (35 mg/mL, 100 mM Tris, pH 8.0) with 0.3 μL of the reservoir solution containing 2 M ammonium sulfate, 100 mM Tris, pH 8,5. All drops were prepared by the crystallization robot Oryx8 (Douglas Instruments, Hungerford, UK) in 96 well 2-lens MRC plates (SWISSCI AG, Neuheim, Switzerland). Crystals reached their maximal size after approximatively one week. To ensure phasing, osmium (IV) hexa-chloride di-potassium salt was added to crystal-containing drops (10 mM final concentration) and left for 24 h. Crystals were then mounted on nylon loops and quickly back-soaked into their reservoir solution + 20% glycerol and immediately flash frozen in liquid nitrogen to be stored until data collection.

All X-ray diffraction experiments were carried out at Synchrotron SOLEIL (St-Aubin, France) on Proxima-2A beamline (Table S1). Measurements were performed at 100 K. A first crystal without osmium was used to collect a native dataset at *λ* = 0.9801 Å with a useful resolution limit of 1.75 Å (data processed with XDS [44] and scaled with Aimless [45]). From an osmium containing crystal, 8 datasets (400° span, 4000 images each) were collected at osmium peak absorption *λ* = 1.1397 Å, at different kappa angles. Merging all of them was mandatory to get a final unique dataset at 2.2 Å resolution with sufficient quality for successful phasing (data processed with XDS and XSCALE). Preliminary phases were determined using the SAD Pipeline implemented in PHASER CCP4i [46] and the partial results were enhanced by ARP/wARP [47]. The resulting model was further improved with the native dataset at 1.75 Å, using PHENIX [48] and COOT [49] (Table S1). The asymmetric unit contained a single OrfG_64-204_ monomer. Several regions were unobserved in the electron density when contoured at 1.2 σ, especially at the N-terminal end. These regions were assumed to be disordered and were not included in the model. The final model contained residues Asp92-Val133 + Val139-Ile168 + Glu176-Ala204, i.e. 70 % of the 143 amino acids of the OrfG_64-204_ subdomain. It is worth noting an important blob of density in the pocket delineated by Tyr106-Phe107, Arg140-Ser143, Tyr164, Glu166 and Phe202 (Fig. S2). This blub could be assigned to none of the compounds used for protein crystallization. It probably resulted from a significant affinity for a ligand present in the protein expression cells or in the successive media used to prepare the protein sample. The final OrfG_64-204_ structure was deposited in the Protein Data Bank with ID 6zgn.

All three-dimensional structure representations were obtained using PYMOL (Schrödinger, LLC).

### Sequence alignment, structure comparison and interface analysis

Protein sequences were retrieved from NCBI and compared using clustalW from MEGA7 [50]. Phylogenetic trees were built with MEGA7 by using the Maximum Likelihood method based on the Tamura-Nei model [51] without outgroup.

The first step of structure comparison was conducted with both DALI [30] and PDBefold [31] to fetch a complete list of VirB8 structures (26 PDB entries). From them, a non-redundant set of 16 structures was conserved (2cc3, 3ub1, 3wz3, 3wz4, 4akz, 4ec6, 4jf8, 4kz1, 4lso, 4mei, 4nhf, 4o3v, 5aiw, 5cnl, 5i97, 6iqt) and added to the new OrfG_64-204_. Then the mTM-align server [52] based on the robust TM-align method [53] performed the structural alignment and produced the structure-based “phylogenetic tree”. PyMOL and an associated homemade python script were used to emphasize visually structure differences and similarities (available upon request). Briefly, a color gradient was applied to each position depending on the rmsd value calculated on Cα carbons of structurally aligned amino acids at this position. Conserved or similar residues were also automatically highlighted. Furthermore, the superposition of all VirB8 monomers produced by the mTM-align server was used to compare oligomers interfaces and to visually identify interesting features.

### Biophysical characterization of native and truncated OrfG_64-331_ proteins

To determine the molecular weight of the purified proteins, size exclusion chromatography (SEC) coupled to MALS analysis was performed on MiniDAWN TREOS II coupled to S200 10/300 increase column (GE Healthcare) mounted on FPLC system (AKTA purifier). We used Tris-HCl 20 mM, pH8, NaCl 100 mM buffer at 0.5 mL/min flow.

### *In vitro* chemical crosslinking

Crosslinking experiments were performed for OrfG_64-331_, OrfG_64-204_ and OrfG_223-331_ obtained after SEC purification and present in Hepes 20 mM pH 8.0, NaCl 100 mM buffer. The used protocol is described as follow: 20 μM of each protein were mixed or not with increasing concentration of paraformaldehyde (PFA) from 0,05 to 5 % in incubated for 1 h at 37°C. The crosslinking reaction was stopped by the adding of Tris 1 M at a final concentration of 80 mM and the mixture was incubated for 10 min at RT. To examine crosslinked products, 15 μL of SDS-PAGE loading buffer (4X) were added and 10 μL from each reaction was analyzed by 12% SDS-PAGE for OrfG_64-331_ and 15% SDS-PAGE for OrfG_64-204_ and OrfG_223-331_.

## Acknowledgments

We thank the members of the DynAMic and CRM2 laboratories for insightful discussions; Laurence Hotel, Emilie Piotrowski, Louise Thiriet, Stéphane Bertin, Johan Staub and Anthony Gauthier for technical assistance; Nicolas Soler and Yvonne Roussel for valuable discussions; Emilie Robert and Jean-Michel Girardet from ASIA platform for encouragements. SEC-MALS analysis were conducted using MiniDAWN TREOS II mounted on FPLC system (AKTA purifier) available on the ASIA platform (Université de Lorraine-INRAE, https://a2f.univ-lorraine.fr/asia/). The Soleil Synchrotron radiation facility is acknowledged for beamline allocation. This work was funded by the National Research Institute for Agriculture, Food and Environment INRAE, the Université de Lorraine and the A2F scientific pole. The project is co-financed by the European Union through the Regional Operational Program of the European Regional Development Fund (ERDF) 2014-2020.

## Authors contribution

JC, AMA, SM, TD, CDJ, BD performed the experiments. NLB: phylogeny analysis. JC developed the script used for structural analysis. CD, FF, NLB, BD: supervision, validation, investigation, methodology. FF, BD: Writing original draft. All authors: review and editing of the manuscript.

